# Tonic activity in lateral habenula neurons promotes disengagement from reward-seeking behavior

**DOI:** 10.1101/2021.01.15.426914

**Authors:** Brianna J. Sleezer, Ryan J. Post, David A. Bulkin, R. Becket Ebitz, Vladlena Lee, Kasey Han, Melissa R. Warden

## Abstract

Survival requires both the ability to persistently pursue goals and the ability to determine when it is time to stop, an adaptive balance of perseverance and disengagement. Neural activity in the lateral habenula (LHb) has been linked to aversion and negative valence, but its role in regulating the balance between reward-seeking and disengaged behavioral states remains unclear. Here, we show that LHb neural activity is tonically elevated during minutes-long disengagements from reward-seeking behavior, whether due to repeated reward omission or following sufficient consumption of reward. Further, we show that LHb inhibition extends ongoing reward-seeking behavioral states but does not prompt re-engagement. We find no evidence for similar tonic activity fluctuations in ventral tegmental area (VTA) dopamine neurons. Our findings implicate the LHb as a key mediator of disengagement from reward-seeking behavior in multiple contexts and argue against the idea that the LHb contributes to decisions solely by signaling aversion.

## INTRODUCTION

Animals transition between directed pursuit of rewards and exploratory or quiescent behavioral states on a timescale of minutes to hours (Ferster and Skinner, 1957; Cohen et al., 2007; Flavell et al., 2013; Hills et al., 2015; Stern et al., 2017; Ebitz et al., 2018; Marques et al., 2020). Factors that influence the persistence of reward-seeking behavioral states include current and predicted homeostatic need (Aponte et al., 2011; Chen et al., 2015), reward proximity (Howe et al., 2013; McGinty et al., 2013; Westbrook and Frank, 2018; Guru et al., 2020), the history of action successes and failures (Vroom, 1964; Charnov, 1976; Ullsperger and von Cramon, 2003; Ebitz et al., 2019), opportunity costs (Niv et al., 2007; Kurzban et al., 2013; Boureau et al., 2015), and environmental threats (Lecca et al., 2017; Alhadeff et al., 2018).

The past decade has seen an intense surge of interest in the role of the lateral habenula (LHb) in regulating reward-seeking behavior. The LHb, part of the epithalamus, is a major conduit of information from the forebrain to brainstem neuromodulatory centers, and regulates behavior at a range of timescales (Bianco and Wilson, 2009; Hikosaka, 2010; Proulx et al., 2014; Hu et al., 2020). At a sub-second timescale, LHb neurons fire phasically when predicted reward is omitted, when cues that predict reward omission or punishment appear, and when shocks or air-puffs are delivered (Matsumoto and Hikosaka, 2007, 2009; Lecca et al., 2017). At longer timescales, neural and metabolic activity in the LHb is elevated during helplessness and passive coping (Caldecott-Hazard et al., 1988; Morris et al., 1999; Shumake et al., 2003; Mirrione et al., 2014; Proulx et al., 2018; Yang et al., 2018; Andalman et al., 2019), and excitatory synaptic transmission onto LHb neurons is potentiated in depression-like behavioral states (Li et al., 2011, 2013; Shabel et al., 2014; Lecca et al., 2016).

Although these findings have been interpreted as evidence that elevated LHb neural activity reflects aversion (Friedman et al., 2011; Stamatakis and Stuber, 2012; Lammel et al., 2012; Proulx et al., 2014), they also raise the possibility that LHb neural activity may simply function as a valence-neutral brake on reward-seeking behavior. Indeed, there is anatomical and behavioral evidence that supports this idea. Midbrain dopamine (DA) neurons play an essential role in supporting sustained goal-directed behavior (Salamone and Correa, 2012; Dolan and Dayan, 2013; Howe et al., 2013; Guru et al., 2020), and the LHb inhibits DA neural activity via the GABAergic rostromedial tegmental nucleus (RMTg) (Christoph et al., 1986; Ji and Shepard, 2007; Matsumoto and Hikosaka, 2007; Hong et al., 2011). LHb lesions reduce reward omission dips in DA neural activity (Tian and Uchida, 2015), LHb stimulation reduces the number of actions that animals are willing to perform for rewards (Proulx et al., 2018), and LHb stimulation reduces entries to spatial locations where stimulation is delivered (Stamatakis and Stuber, 2012). Intriguingly, lesioning the LHb or silencing excitatory inputs to the LHb elevates consumption of palatable food but not contaminated food (Paul et al., 2011; Stamatakis et al., 2016). These results are consistent with a role for the LHb in suppressing reward-seeking behavior but more difficult to reconcile with a model in which the primary role of the LHb is to signal aversion.

Here, we combine fiber photometry, optogenetics, and multi-unit electrophysiology in freely behaving mice to investigate the role of tonic LHb neural activity in regulating the persistence of reward-seeking behavioral states. Our findings reveal that large, sustained increases in tonic LHb neural activity accompany minutes-long disengagements from reward-seeking behavior, and that these tonically excited LHb states can be driven either by task disengagement due to repeated reward omission (negatively valenced state) or by spontaneous task disengagement following the consumption of sufficient reward (positively valenced state). Further, we demonstrate a causal role for LHb neural activity in regulating reward-seeking behavior by showing that LHb inhibition extends reward-seeking behavioral states. Our results strongly support the hypothesis that a key role of tonic LHb neural activity is to promote disengagement from reward-seeking behavior.

## RESULTS

### Calcium Dynamics in Genetically Targeted LHb Neurons

The LHb and surrounding brain tissue are composed primarily of glutamatergic neurons, and achieving expression of genetically encoded tools in the LHb while confidently excluding expression in adjacent brain regions is challenging. In the Klk8-Cre (NP171) mouse line Cre recombinase is expressed in the habenula but not in adjacent brain regions (GENSAT; Proulx et al., 2014), a feature that allowed us to restrict the expression of genetically-encoded optical indicators and actuators. Although Cre is also expressed in the medial habenula (MHb) in this line, we were able to consistently minimize MHb expression by refinement of viral vector serotype, injection volume, and coordinates (see Methods). To characterize Cre-dependent gene expression in Klk8-Cre (NP171) mice, we injected AAV9-CAG-Flex-GFP in the LHb and quantified co-localization of GFP and NeuN, a marker of neuronal identity (Mullen et al., 1992). We found that GFP expression was highly specific to neurons (97.8% +/- 0.49% specificity), and that most NeuN-labeled neurons at the injection site were GFP-positive (71.6% +/- 2.97% penetrance, **Figure 1A**).

**Figure 1.**
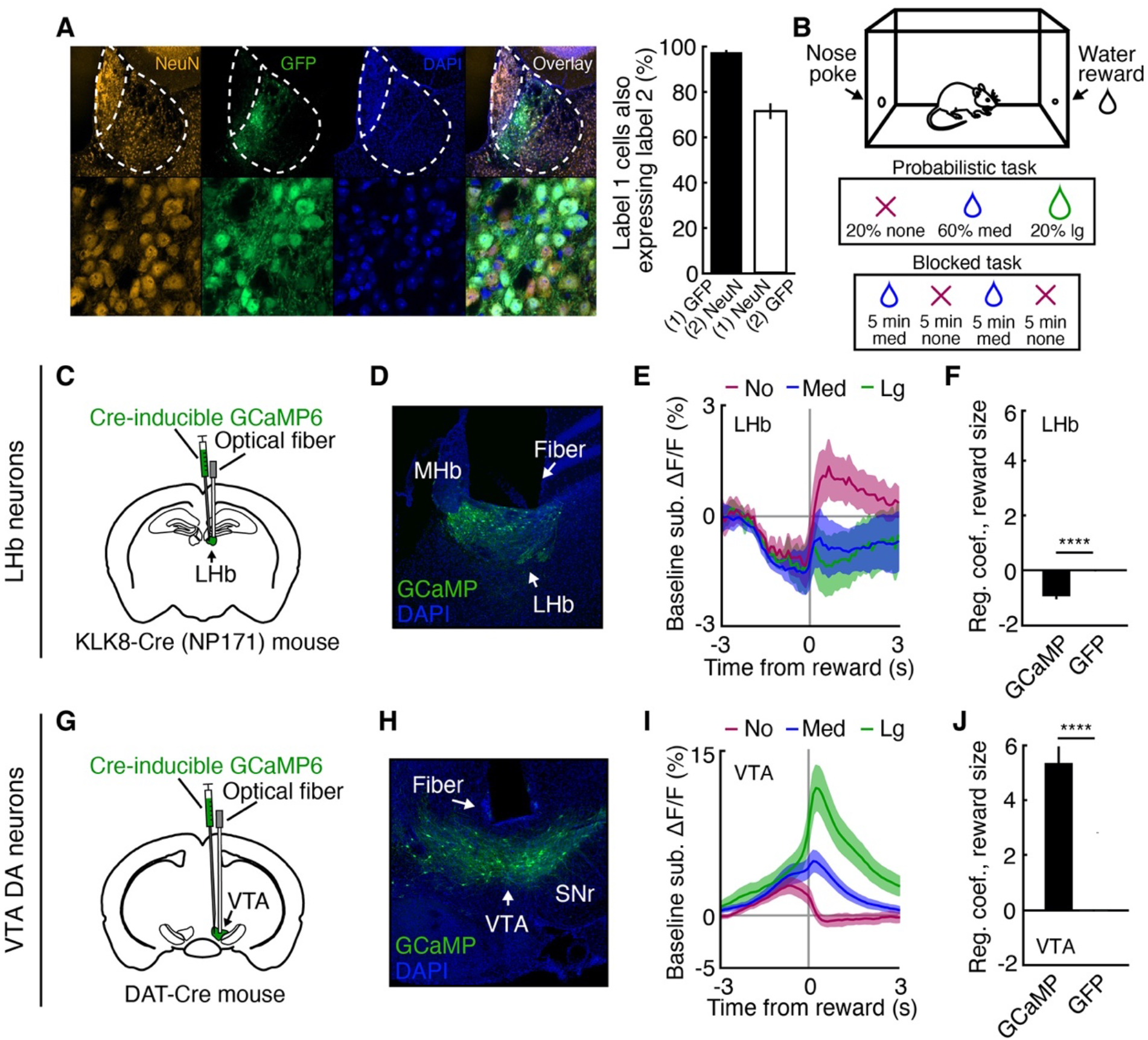
Calcium Dynamics in Genetically Targeted LHb Neurons. (A) Colocalization of NeuN and GFP expression in the LHb of Klk8-Cre (NP171) mice. (B) Schematic of probabilistic and blocked poke-reward task designs. (C) Schematic of LHb viral vector injection and optical fiber placement. (D) GCaMP6 expression in LHb neurons in a Klk8-Cre (NP171) mouse. (E) Average baseline-subtracted ΔF/F in GCaMP Klk8-Cre (NP171) mice (n = 7) during probabilistic task sessions, aligned to reward receipt and separated by reward size (no reward, red; medium reward, blue; large reward, green). (F) Average regression coefficients obtained by regressing ΔF/F 0-1 s following the start of reward consumption onto reward size in GCaMP (n = 7) and GFP (n = 3) Klk8-Cre (NP171) mice. (G-J) Same as C-F, but for VTA DA neurons in GCaMP (n = 6) and GFP (n = 3) DAT-Cre mice. ****p < 0.0001, two-sample t-test. Shaded regions and error bars indicate SEM. See also Figure S1.

To further validate use of the Klk8-Cre (NP171) mouse line for LHb neural targeting, we used fiber photometry to monitor LHb population neural dynamics during reward delivery and omission (Cui et al., 2013; Gunaydin et al., 2014). We injected a genetically-encoded calcium indicator, AAV9-CAG-Flex-GCaMP6s (Chen et al., 2013), into the LHb of Klk8-Cre (NP171) mice, and implanted an optical fiber over the LHb to monitor calcium-dependent fluorescence (**Figures 1C and 1D**). Because LHb neurons are inhibited by reward delivery and excited by the omission of predicted rewards (Matsumoto and Hikosaka, 2007, 2009; Bromberg-Martin and Hikosaka, 2011), we examined LHb activity during performance of a simple self-paced operant task. In this ‘poke-reward’ task, mice poked their nose into a port on one side of an operant chamber in order to trigger the delivery of a water reward on the other side (**Figure 1B**). We recorded LHb activity while mice performed a probabilistic version of this task in which each trial had a 20% chance to yield no reward, a 60% chance to yield a medium reward (10 µl), and a 20% chance to yield a large reward (20 µl). As expected, LHb neural activity was negatively correlated with reward size (GCaMP6s (n = 7), GFP (n = 3), p < 0.0001, two-sample t-test, **Figures 1E and 1F**).

For validation, we recorded LHb activity in Vglut2-ires-Cre mice (Vong et al., 2011) using fiber photometry (**Figures S1A and S1B)**, and in C57BL6/J mice using multi-unit electrophysiology (**Figures S1E and S1F**) during performance of the same task. Both LHb Vglut2 neural activity (GCaMP6 (n = 3), GFP (n = 3), p < 0.0001, two-sample t-test, **Figures S1C and S1D**) and LHb multi-unit activity (n = 40 electrodes in 6 mice, p < 0.0001, two-sample t-test, **Figures S1G and S1H**) were negatively correlated with reward size, consistent with our findings in Klk8-Cre (NP171) mice and further supporting the use of these mice for investigating LHb neural dynamics and function. We also recorded activity from VTA DA neurons in DAT-Cre mice (Bäckman et al., 2006) during performance of the same task. Consistent with previous findings (Schultz et al., 1997; Matsumoto and Hikosaka, 2007; Cohen et al., 2012), we found that VTA DA neurons were phasically excited during delivery of medium and large rewards and inhibited during reward omission (GCaMP6 (n = 6), GFP (n = 3), p < 0.0001, two-sample t-test, **Figures 1G-1J**).

### Tonically Elevated LHb Activity During Spontaneous End-of-Session Task Disengagement

Intriguingly, the largest LHb activity changes we observed were periods of sustained excitation that occurred toward the end of poke-reward task sessions when mice began to spontaneously disengage from task performance (**Figures 2A and 2B**). LHb neural activity was tonically elevated throughout these disengaged periods and decreased if mice spontaneously re-engaged in the task. To quantify this effect, we used a two-state hidden Markov model (HMM, see Methods) to identify periods of time during which mice were less engaged in task performance (low task engagement states) and periods during which mice were more strongly engaged (high task engagement states). We have previously used this approach to differentiate periods of exploration and exploitation (Ebitz et al., 2018, 2019). We regressed task engagement state onto ΔF/F for each session, and included running speed as a predictor to assess whether locomotor activity may also contribute to tonic LHb activity. LHb neural activity was negatively correlated with task engagement state, but was not correlated with either running speed or the interaction between task engagement state and running speed (GCaMP6 (n = 7), GFP (n = 3), engagement state: p < 0.0001, running speed: p = 0.7287, engagement state X running speed: p = 0.9637, two-sample t-test, **Figure 2C**). Thus, tonically elevated LHb neural activity reflects spontaneous disengagement from reward-seeking task performance.

**Figure 2.**
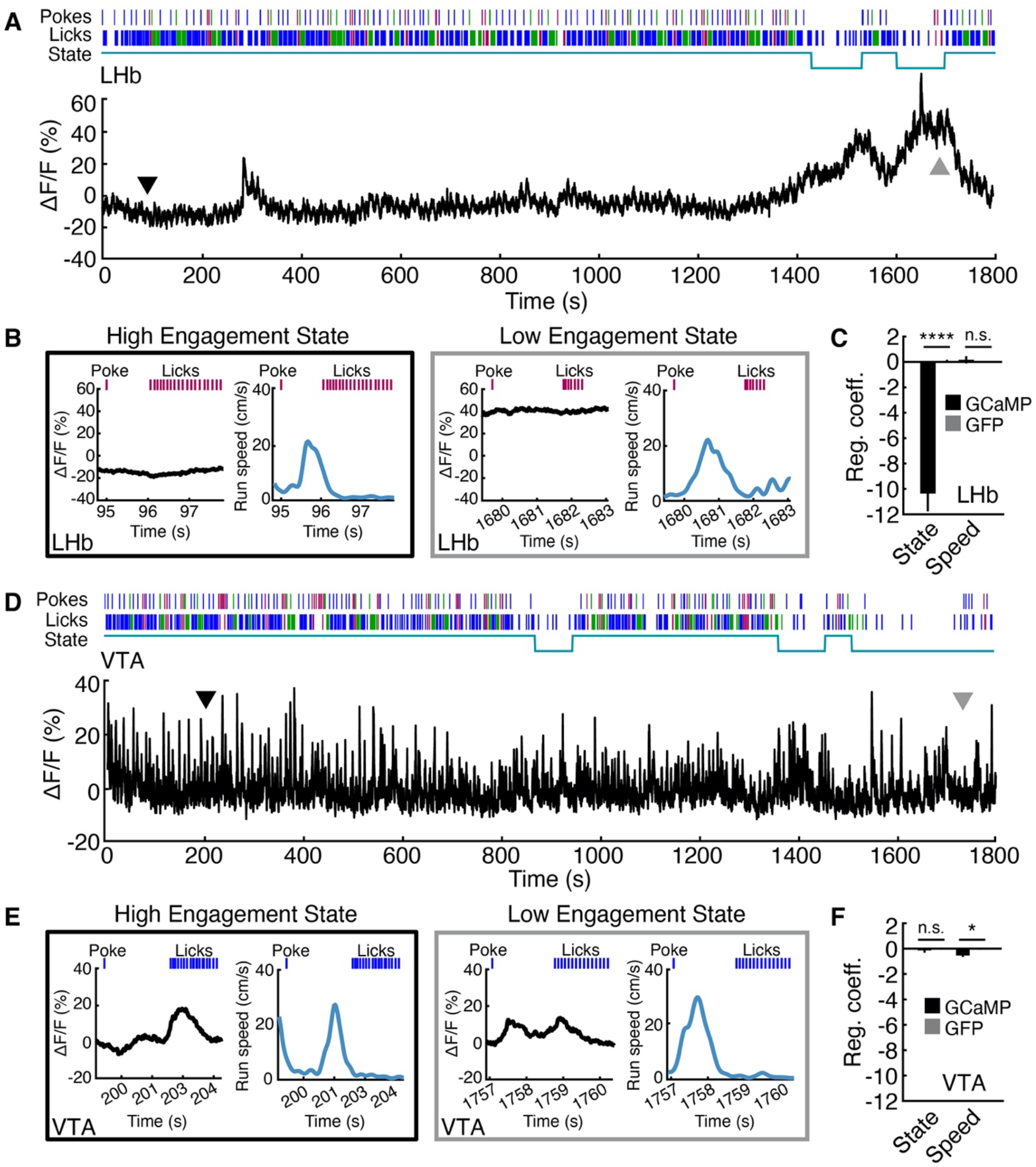
Tonically Elevated LHb Activity During Spontaneous End-of-Session Task Disengagement. (A) Example LHb Klk8-Cre (NP171) photometry from a complete probabilistic task session. Lines above the plot indicate all poke and lick times (no reward, red; medium reward, blue; large reward, green). The line under the poke and lick timestamps indicates whether mice were in a high or low task engagement state. Black and gray arrows correspond to example trials shown in B. (B) Two single trials from the example photometry trace in A. Left: trial during high engagement state. Right: trial during low engagement state. (C) Average regression coefficients obtained by regressing ΔF/F from each session onto engagement state and running speed in GCaMP (n = 7) and GFP (n = 3) Klk8-Cre (NP171) mice. (D) Example VTA DAT-Cre photometry from a complete probabilistic task session, same conventions as in A. Black and gray arrows correspond to example trials shown in E. (E) Two single trials from the example photometry trace in D. Left: trial during high engagement state. Right: trial during low engagement state. (F) Average regression coefficients obtained by regressing ΔF/F onto engagement state and running speed in GCaMP (n = 6) and GFP (n = 3) DAT-Cre mice. *p < 0.05, ****p < 0.0001, two-sample t-test. Error bars indicate SEM. See also Figure S2.

Findings from Vglut2-Cre mice were similar (GCaMP6 (n = 3), GFP (n = 3), engagement state: p = 0.0029, running speed: p = 0. 2454, two-sample t-test, **Figures S2A and S2B**), but we observed a positive correlation between LHb neural activity and speed in multi-unit electrophysiology data (n = 40 electrodes in 6 mice, engagement state: p < 0.0001, running speed: p < 0.0001, t-test compared to shuffled sample, **Figures S2C and S2D**). Interestingly, when we examined VTA DA neural activity in DAT-Cre mice during this task we found no correlation between neural activity and engagement state or the interaction between engagement state and running speed (GCaMP6 (n = 6), GFP (n = 3), engagement state: p = 0.6767, engagement state X running speed: p = 0.1865, two-sample t-test, **Figures 2D-2F**), but neural activity was negatively correlated with running speed alone (running speed: p = 0.0203, two-sample t-test). The absence of tonic VTA DA neural activity changes between high and low task engagement states is reminiscent of previous reports of stable baseline spiking activity in VTA DA neurons through changing schedules of rewards or punishments (Cohen et al., 2015; Mohebi et al., 2019).

### Baseline Tonic LHb Activity Reflects Task Engagement State

To confirm that tonic disengagement-related LHb excitation was not simply a consequence of differences in phasic reward-related activity across high and low engagement states, we examined average LHb activity during two specific task epochs in both high and low engagement states: the baseline epoch (1s prior to each trial-initiating poke) and the reward epoch (1s following the first lick). We found greater mean LHb neural activity in low compared to high engagement states during both epochs in Klk8-Cre (NP171) mice (n = 7, baseline: p = 0.0003, reward: p = 0.0001, paired samples t-test, **Figures 3A and 3B**). We found the same effect in Vglut2-Cre mice (n = 3, baseline: p = 0.0115, reward: p = 0.0104, paired samples t-test, **Figures S3A and S3B**) and via multi-unit electrophysiology (n = 40 electrodes in 6 mice, baseline: p < 0.0001, reward: p < 0.0001, paired samples t-test, **Figures S3C and S3D**). In contrast, VTA DA neural activity during the baseline epoch did not depend on task engagement state (n = 6, p = 0.4183, paired samples t-test, **Figures 3C and 3D**). However, VTA DA neural activity during the reward epoch was significantly greater in high engagement states (n = 6, p < 0.0001, paired samples t-test, **Figures 3C and 3D**), likely due to the enhanced phasic VTA DA reward response in this state (Bassareo and Chiara, 1999; Branch et al., 2013; Papageorgiou et al., 2016).

**Figure 3.**
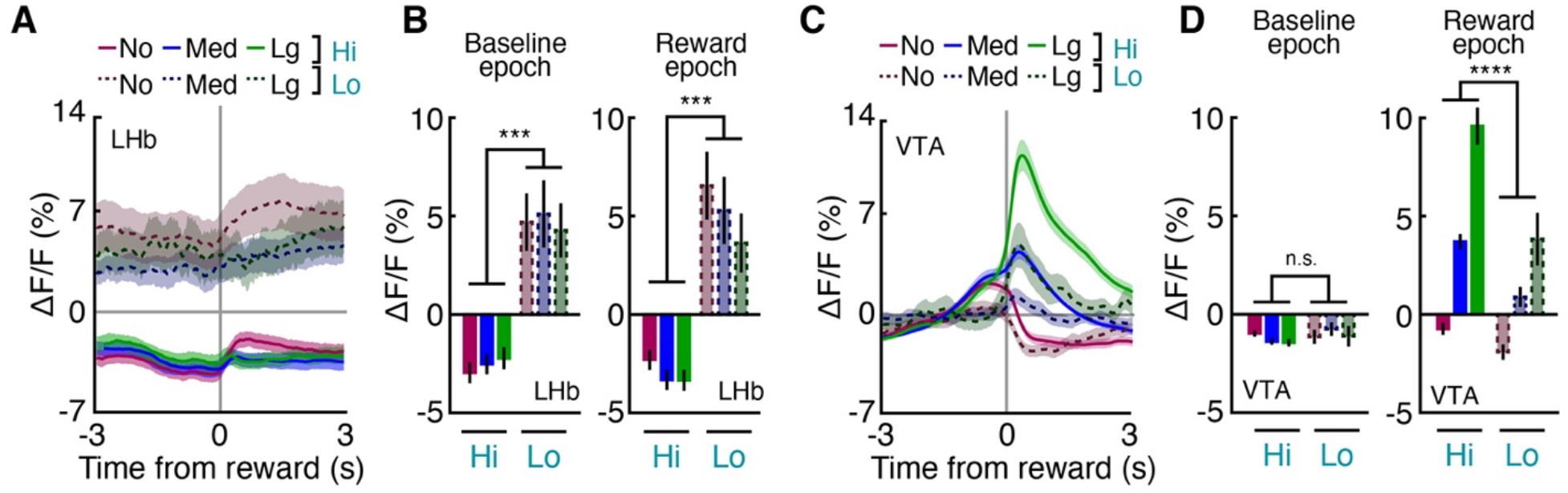
Baseline LHb Activity Reflects Task Engagement State. (A) Average ΔF/F in GCaMP Klk8-Cre (NP171) mice (n = 7) during probabilistic task sessions, aligned to reward receipt and separated by engagement state (solid versus dotted lines) and reward size (no reward, red line; medium reward, blue; large reward, green). (B) Average ΔF/F during baseline and reward epochs in high and low engagement states. (C-D) Same as A-B but for VTA DA neurons in DAT-Cre mice (n = 6). ***p < 0.001, ****p < 0.0001, paired samples t-test. Error bars and shaded regions indicate SEM. See also Figure S3.

Task disengagements occurring near the end of each 30-minute task session suggest satiety, but it is also possible that disengagements occurred because of fatigue or reward omission. If fatigue induced task disengagement, mice might be expected to move less during low engagement states. To investigate this possibility we compared locomotor activity during periods of high and low task engagement, but we found that average speed was not significantly different between states (high engagement speed: 5.6326 ± 0.1348 cm/s, low engagement speed: 5.0406 ± 0.1062 cm/s, n = 105 sessions, p = 0.6031, two-sample t-test). If reward omission drove disengagement, entries into low engagement states would be preceded by non-rewarded trials more often than expected by chance. Contrary to this hypothesis, we found that entries into low engagement states were actually less likely than chance to be preceded by reward omissions and more likely than chance to be preceded by medium or large rewards (no reward: p < 0.0001, medium reward: p = 0.0037, large reward: p = 0.0001, binomial test, **Figure S3E**). Thus, mice were more likely to disengage after rewards that brought them closer to satiety. We also found that entries into high engagement states were more likely than chance to be preceded by reward omissions and less likely than chance to be preceded by medium rewards (no reward: p < 0.0001, medium reward: p < 0.0001, large reward: 0.8418, binomial test, **Figure S3E**), which may reflect the extinction burst (Skinner, 1938; Cooper et al., 1987). Together, these findings argue against the hypothesis that fatigue or reward omission drove entry into low engagement states in the probabilistic poke-reward task and suggest that mice disengaged from task performance toward the end of sessions because they had satisfied their homeostatic need.

### LHb Activity is Tonically Elevated During Task Disengagement Due to Repeated Reward Omission

We next asked whether tonic LHb excitation is specific to end-of-session task disengagement, or if LHb neural activity might also be elevated when mice disengage in response to other factors such as repeated reward omission. To address this question, we recorded LHb neural activity in Klk8-Cre (NP171) mice while mice performed a blocked version of the poke-reward task in which 5-minute blocks of medium-reward trials alternated with 5-minute blocks of no-reward trials (**Figure 1B**). As in the probabilistic poke-reward task, we found that LHb neural activity was high during task disengagements and low during on-task behavior (**Figures 4A and 4B**). To quantify this effect, we regressed task engagement state onto ΔF/F for each session and included block reward contingencies and running speed as predictors. LHb neural activity was negatively correlated with task engagement state and block reward, but was not correlated with running speed (GCaMP6 (n = 6), GFP (n = 3), engagement state: p = 0.0295, block reward: p = 0.0268, running speed: p = 0.4364, two-sample t-test, **Figure 4C**). Regression coefficients for two- and three-way interactions were not different between GCaMP and GFP groups (p > 0.05 for all comparisons, paired samples t-tests). These results were confirmed in Vglut2-Cre mice (GCaMP6 (n = 3), GFP (n = 3), engagement state: p = 0.0145, block reward: p = 0.6470, running speed: p = 0.1563, two-sample t-test) and via multiunit electrophysiology (n = 40 electrodes in 6 mice, engagement state: p < 0.0001, block reward: p < 0.0001, running speed: p < 0.0001, two-sample t-test, **Figure S4**). VTA DA neural activity in DAT-Cre mice positively correlated with block reward, but we found no correlation with engagement state or running speed (GCaMP6 (n = 6), GFP (n = 3), engagement state: p = 0.7294, block reward: p = 0.0407, running speed: p = 0.5685, two-sample t-test, **Figures 4D-4F**). Together, these data reveal that tonically elevated activity in LHb neurons accompanies task disengagements prompted by factors with both negative and positive valence.

**Figure 4.**
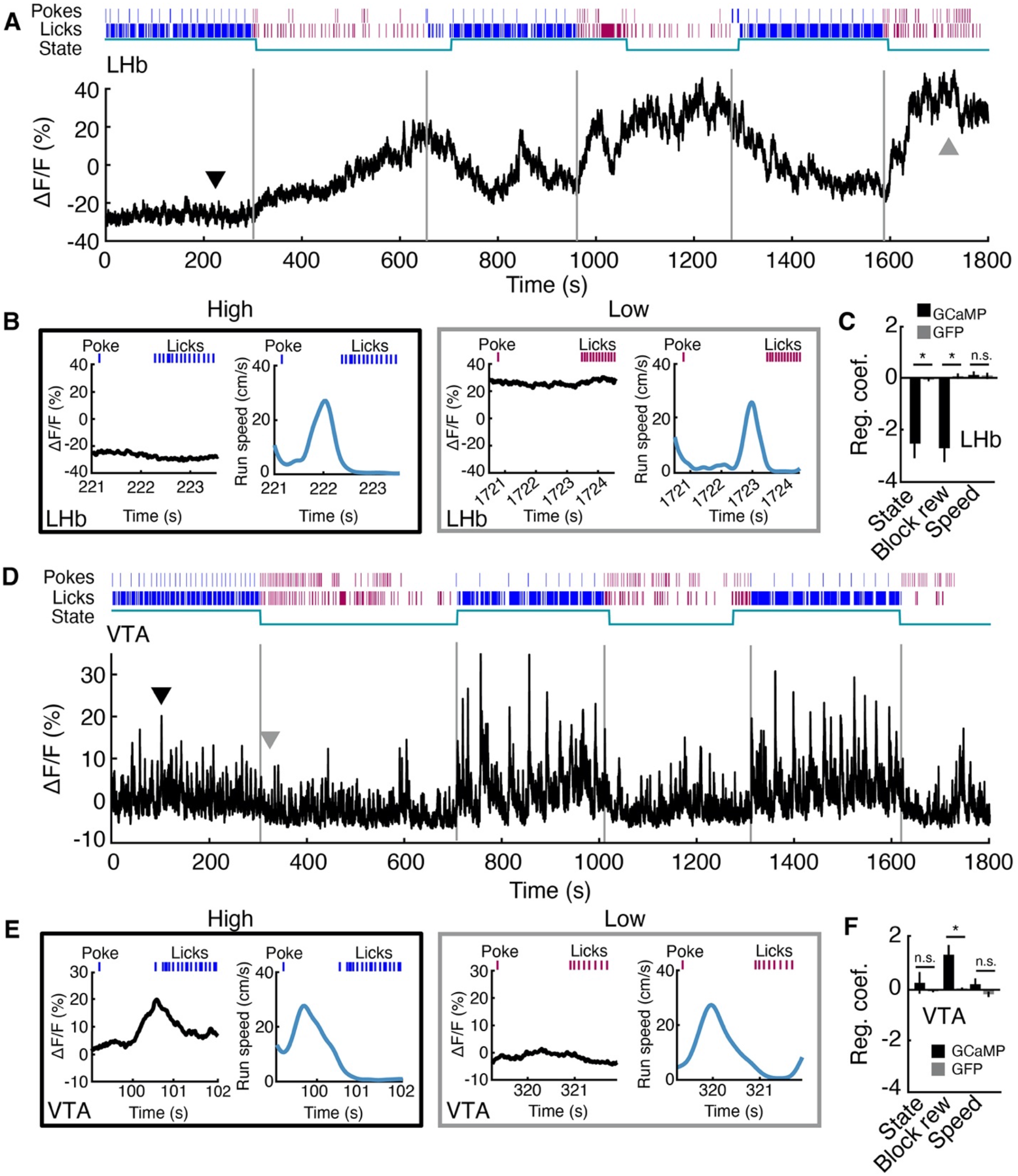
LHb Activity is Tonically Elevated During Task Disengagement Due to Repeated Reward Omission. (A) Example LHb Klk8-Cre (NP171) photometry during a blocked task session. Lines above the plot indicate all poke and lick times (no reward, red; medium reward, blue). The line under the poke and lick timestamps indicates whether mice were in a high or low task engagement state. Black and gray arrows correspond to example trials shown in B. (B) Two single trials from the example photometry trace in A. Left: trial during high engagement state. Right: trial during low engagement state. (C) Average regression coefficients obtained by regressing ΔF/F onto engagement state, block reward, and running speed in GCaMP (n = 6) and GFP (n = 3) Klk8-Cre (NP171) mice. (D) Example VTA DAT-Cre photometry during a blocked task session, same conventions as A. Black and gray arrows correspond to example trials shown in E. (E) Two single trials from the example photometry trace in D. Left: trial during high engagement state. Right: trial during low engagement state. (F) Average regression coefficients obtained by regressing ΔF/F onto engagement state, block reward, and running speed in GCaMP (n = 6) and GFP (n = 3) DAT-Cre mice. *p < 0.05, two-sample t-test. Error bars indicate SEM. See also Figure S4.

### Rising LHb Activity Precedes Task Disengagement

In order to characterize the emergence of elevated tonic LHb activity during task disengagement, we examined the trials leading up to and following transitions into low engagement states. We averaged LHb neural activity during the baseline period (1 s prior to the trial-initiating poke) for each of the 10 trials preceding disengagement, and we calculated the slope of the change in average activity from the first to the last of these trials. In the probabilistic poke-reward task, we observed that average LHb activity in Klk8-Cre (NP171) mice increased across the trials preceding disengagement, while average VTA DA activity did not (Klk8-Cre: GCaMP6 (n = 7), GFP (n = 3), DAT-Cre: GCaMP6 (n = 6), GFP (n = 3), Klk8-Cre slope: p = 0.0195, DAT-Cre slope: p = 0.4850, two-sample t-test, **Figures 5A and 5B**). We found similar effects during the blocked poke-reward task (Klk8-Cre: GCaMP6 (n = 6), GFP (n = 3), DAT-Cre: GCaMP6 (n = 6), GFP (n = 3), Klk8-Cre slope: p = 0.0060, DAT-Cre slope: p = 0.7910, two-sample t-test, **Figures 5C and 5D**). To determine how far in advance of the state transition LHb tonic activity began to increase, we calculated the slope of the change in baseline activity across a symmetrical 10 trial window slid in 1 trial steps across the transition into low engagement states. For both tasks, LHb activity began increasing about 10 trials prior to entry into low engagement states (**Figure 5E-F**).

**Figure 5.**
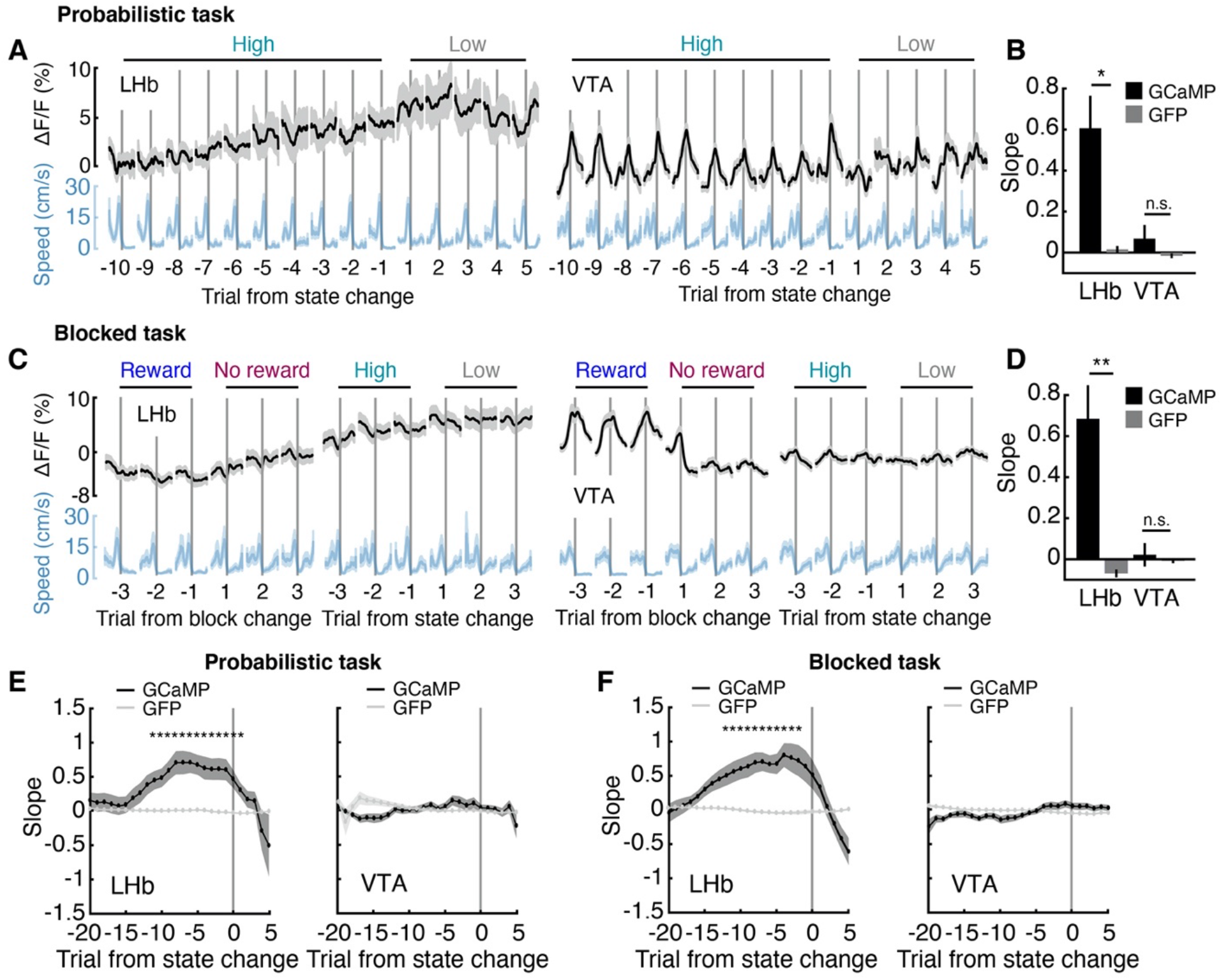
Rising LHb Activity Precedes Task Disengagement. (A) Average reward-aligned ΔF/F for the trials immediately before and after entry into low engagement states, probabilistic task sessions. Left, LHb ΔF/F in Klk8-Cre (NP171) mice. Right: VTA DA ΔF/F. (B) Slope of the increase in baseline ΔF/F over the 10 trials before entry into low engagement states in LHb Klk8-Cre (NP171) (GCaMP (n = 7), GFP (n = 3)) and VTA DAT-Cre (GCaMP (n = 6), GFP (n = 3)), probabilistic task sessions. (C) Average reward-aligned ΔF/F for the trials immediately before and after the start of no-reward blocks and entry into low engagement states, blocked task sessions. Left, LHb ΔF/F in Klk8-Cre (NP171) mice. Right: VTA DAT-Cre ΔF/F. (D) Slope of the increase in baseline ΔF/F over the 10 trials before entry into low engagement states in LHb (GCaMP (n = 6), GFP (n = 3)) and VTA (GCaMP (n = 6), GFP (n = 3)), blocked task sessions. (E) Slope of the increase in baseline ΔF/F over a 10 trial sliding window, probabilistic task. (F) Slope of the increase in baseline ΔF/F over a 10 trial sliding window, blocked task. *p < 0.05, **p < 0.01, two-sample t-test. Error bars indicate SEM.

### LHb Inhibition Extends Reward-Seeking Behavioral States

If tonically elevated LHb neural activity promotes task disengagement, inhibiting the LHb should disrupt the ability to disengage from on-task behavior. To test this hypothesis, we optogenetically inhibited LHb neurons while mice performed the poke-reward task. We bilaterally injected AAV-EF1α-DIO-eNpHR3.0-eYFP into the LHb of Klk8-Cre (NP171) mice and implanted optical fibers above the LHb for light delivery (**Figures 6A and 6B**). We asked two questions: 1) does LHb inhibition during disengaged states prompt re-engagement in task performance? 2) does LHb inhibition during task-engaged states disrupt the ability to disengage from task performance?

**Figure 6.**
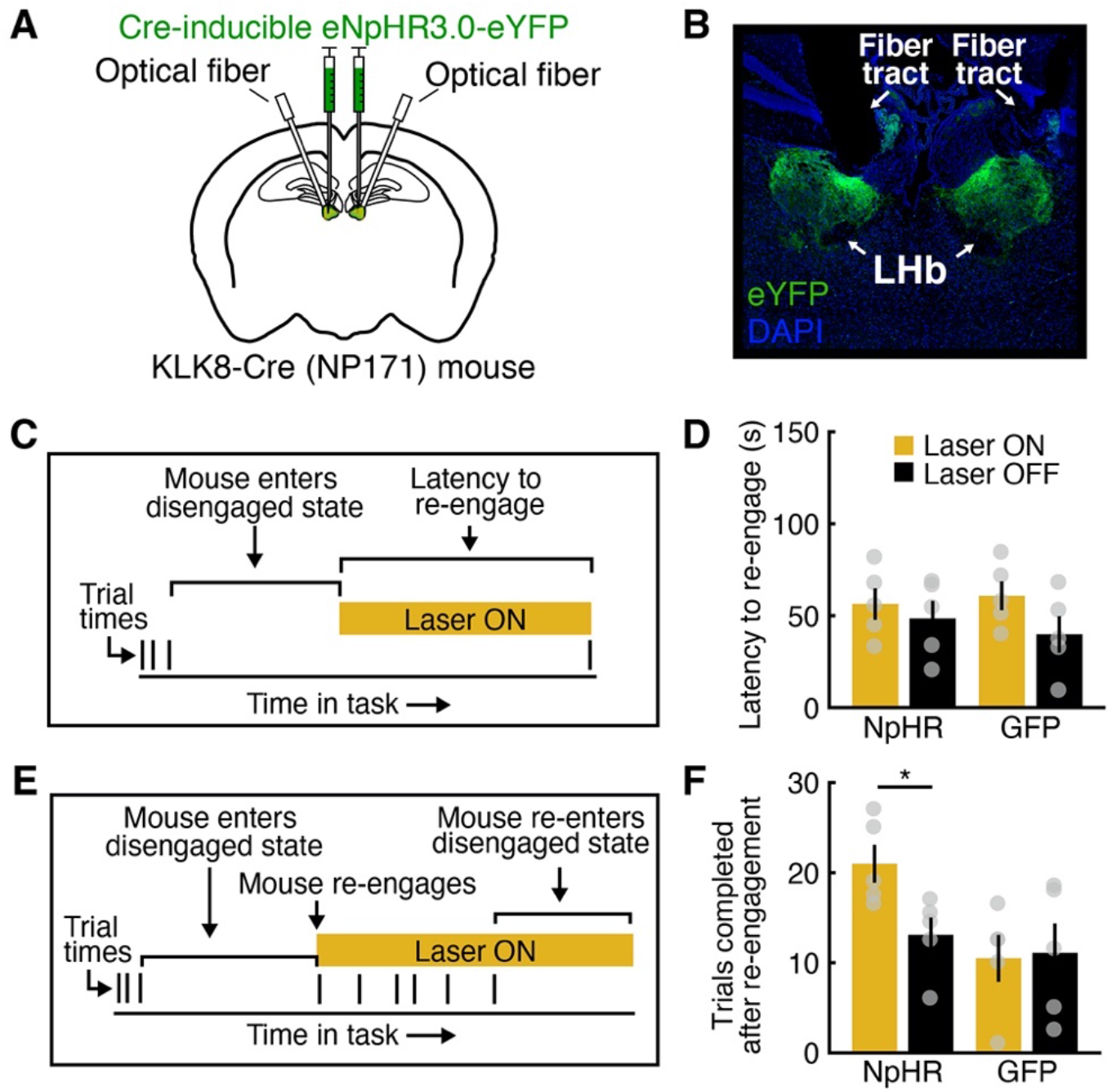
LHb Inhibition Extends Reward-Seeking Behavioral States. (A) Schematic of LHb viral vector injection and optical fiber placement. (B) eNpHR3.0-eYFP expression in LHb neurons in a Klk8-Cre (NP171) mouse. (C) Experimental schematic, task re-engagement. (D) Average latency to re-engage in task performance. LHb inhibition initiated during disengagement. NpHR (n = 5) and GFP (n = 5) Klk8-Cre (NP171) mice. (E) Experimental schematic, task persistence. (F) Average number of trials completed after task re-engagement. LHb inhibition initiated upon re-engagement. NpHR (n = 5) and GFP (n = 5) Klk8-Cre (NP171) mice. *p < 0.05, paired samples t-test. Error bars indicate SEM. See also Figure S5.

To examine whether LHb inhibition during disengaged states prompts task re-engagement, we allowed mice to freely perform poke-reward trials for medium rewards and waited until they spontaneously refrained from task performance for two minutes. At this point light delivery was initiated, and inhibition was maintained until mice spontaneously performed another trial (**Figure 6C**). We found that inhibiting LHb during disengaged states had no effect on the latency to re-engage (NpHR (n = 5), GFP (n = 5), p = 0.6669, paired samples t-test, **Figure 6D**). To examine whether LHb inhibition disrupts the ability to disengage we began as above, but instead of starting inhibition after two minutes without trials we continued to wait until mice spontaneously completed another trial. At this point light delivery was initiated, and inhibition was maintained until mice again refrained from task performance for two minutes. Here, we found that LHb inhibition increased the mean number of trials completed before the next disengagement (NpHR (n = 5), GFP (n = 5), p = 0.0316, paired samples t-test, **Figure 6E**). Thus, LHb inhibition extends task-engaged states but does not prompt re-engagement in disengaged mice.

Finally, we asked whether extended task engagement upon LHb inhibition was specific to conditions in which reward was available, or whether LHb inhibition promoted persistent task performance even in the absence of reward. To address this question, mice were allowed to perform the poke-reward task as above, but no rewards were delivered at any time during task performance during this session. Here, we found no effect of LHb inhibition on either the latency to re-engage (NpHR (n = 5), GFP (n = 5), p = 0.8208, paired samples t-test, **Figure S6A**) or on the number of trials completed before the next disengagement (NpHR (n = 5), GFP (n = 5), p = 0.3367, paired samples t-test, **Figure S6B**). Thus, LHb inhibition only prolongs task-engaged states when rewards are available. Together, these findings support a framework where LHb neural activity serves as a brake on the neural systems that promote reward-seeking behavior.

## DISCUSSION

Here, we sought to investigate how LHb neural activity regulates the balance between engaged reward-seeking and disengaged behavioral states. LHb recordings during a self-paced reward-seeking task revealed large, sustained increases in neural activity that occurred during minutes-long task disengagements. We observed these activity increases both when disengagement followed repeated reward omission and when disengagement occurred spontaneously at the end of task sessions following sufficient reward consumption. Phasic LHb neural activity upon reward omission was also observed, but was moderate in comparison to state-dependent tonic changes. In contrast, we observed large phasic reward signals in VTA DA neural activity but did not detect state-dependent tonic changes, consistent with previous studies (Cohen et al., 2015; Mohebi et al., 2019). Finally, we found that inhibiting LHb neurons prolonged ongoing reward-seeking behavioral states but did not prompt disengaged mice to resume task performance.

Our findings suggest that LHb neural activity may function as a brake or stop signal (Seeley et al., 2012), potentially acting in opponency to DA signals that promote reward-seeking behavior, a canonical circuit architecture that can be found in neural circuits ranging from the retina to essential escape circuits (Joesch and Meister, 2016; Koyama et al., 2016) and which has an extensive theoretical foundation in evidence accumulation models of perceptual decision making (Shadlen and Newsome, 2001; Gold and Shadlen, 2007; Bogacz et al., 2006; van Ravenzwaaij et al., 2012). LHb neural activity has been linked to the cessation of motor output (Hikosaka, 2010), but it also has been shown to facilitate active escape behavior (Lecca et al., 2017), both of which are compatible with a primary role for LHb circuits in prompting disengagement from reward-seeking behavior.

Disengagement from reward-seeking behavior can be triggered by a variety of factors with positive or negative valence. For example, an animal might stop attempting to obtain water either because it has already consumed enough water and its homeostatic needs have been met, or because its actions to obtain water have proven ineffective. Homeostatic resolution is positively valenced (Betley et al., 2015; Garfield et al., 2015; Schéle et al., 2017) and action failure is negatively valenced (Skinner, 1953; Leitenberg, 1965; Amsel, 1992; Papini and Dudley, 1997), but both factors prompt task disengagement. Supporting this notion, the LHb receives its strongest afferents from basal ganglia circuits involved in action selection and evaluation and hypothalamic circuits involved in homeostatic regulation (Tachibana and Hikosaka, 2012; Jennings et al., 2013; Mahler et al., 2014; Root et al., 2015; Stephenson-Jones et al., 2016; Chang et al., 2017; Ottenheimer et al., 2018). Inputs from lateral hypothalamus, lateral preoptic area, ventral pallidum, and entopeduncular nucleus/internal globus pallidus are particularly robust (Herkenham and Nauta, 1977; Hong and Hikosaka, 2008; Shabel et al., 2012, 2014; Proulx et al., 2014; Yetnikoff et al., 2015; Stamatakis et al., 2016; Barker et al., 2017; Lecca et al., 2017; Zahm and Root, 2017; Lazaridis et al., 2019).

Other LHb inputs arise from brainstem regions essential for defensive behavior, such the periaqueductal gray, superior colliculus, and raphe nuclei (Zhao et al., 2014; Shang et al., 2015; Yetnikoff et al., 2015; Tovote et al., 2016; Zahm and Root, 2017; Huang et al., 2017; Evans et al., 2018; Seo et al., 2019). LHb neurons fire when animals receive air puffs and shocks that trigger escape or startle movements (Matsumoto and Hikosaka, 2009; Lecca et al., 2017), and silencing glutamatergic inputs to LHb neurons impairs escape (Lecca et al., 2017). We hypothesize that in threatening situations LHb neural activity may play an essential role in terminating ongoing reward-seeking behavioral states in order to permit defensive circuits to quickly assume control of behavior. The LHb also receives inputs from frontal cortical regions essential for behavioral flexibility (Greatrex and Phillipson, 1982; Miller and Cohen, 2001; Monsell, 2003; Kim and Lee, 2012), and LHb inactivation impairs task switching (Thornton and Evans, 1982; Baker et al., 2015; Baker and Mizumori, 2017). Thus, a diverse array of LHb afferents provides potential mechanistic substrates for disengaging from ongoing reward-seeking behavior in response to a variety of negatively and positively valenced factors.

An intriguing finding was the lack of tonic engagement-related signals we observed in VTA DA neuron activity, particularly given that the LHb is thought to contribute to reward related processing primarily through the inhibition of VTA DA neurons (Christoph et al., 1986; Ji and Shepard, 2007; Matsumoto and Hikosaka, 2007; Balcita-Pedicino et al., 2011; Tian and Uchida, 2015). Our findings suggest that LHb is not simply an inverse of VTA DA neuron activity. Interestingly, the tonic activity fluctuations we observed in LHb resemble fluctuations in striatal DA release that reflect motivational state (Mohebi et al., 2019). If long timescale engagement-related signals are present in tonic LHb activity and striatal DA release, but not VTA DA neuron activity, which downstream circuits might be contributing to the regulation of sustained reward-seeking and disengaged behavioral states? One possibility is that signals related to sustained disengagement are transmitted from LHb to the raphe nuclei, but recent evidence showing that chemogenetic inhibition of the LHb-dorsal raphe projection reduces perseverative reward seeking argues against this hypothesis (Coffey et al., 2020). However, the LHb also sends a major projection to the median raphe which may serve a distinct function (Proulx et al., 2014).

Another interesting finding was the behavioral specificity of LHb inhibition. While LHb inhibition extended task engagement, it did not facilitate task re-engagement. Further, LHb inhibition had no effect on persistence if rewards were not available. These findings are consistent with the idea of LHb as a brake on reward-seeking behavior – releasing the brake would not promote task re-engagement if a signal that promotes reward-seeking behavior is not active.

Our findings strongly support the hypothesis that a key role of tonic LHb neural activity is to promote disengagement from reward-seeking behavior, suggest that tonic LHb neural activity acts as a valence-neutral brake on reward-seeking behavior, and add to mounting evidence that the functional role of LHb neural activity extends beyond the processing of negative events or costs. This shift in our understanding of LHb circuit function is essential for assessing the potential and limitations of the LHb as a therapeutic entry point for mood and anxiety disorders, and provides a conceptual framework of potential utility for deepening our understanding of the functional roles of downstream neuromodulatory circuits.

## ACKNOWLEDGMENTS

We thank I.T. Ellwood, J.R. Fetcho, H.K. Reeve, J.H. Goldberg, A. Guru, C. Seo, E.L. Troconis, Y.Y. Ho, W. Gu, and Y. Baumel for helpful discussions; A.K. Recknagel for expert technical assistance; and the Warden laboratory and Cornell Neurobiology and Behavior for training and support. Supported by the Mong Family Foundation (R.J.P., D.A.B.), the Brain and Behavior Research Foundation (D.A.B., M.R.W.), the New York Stem Cell Foundation (M.R.W.), an NIH DP2 New Innovator Award (M.R.W.), the Alfred P. Sloan Foundation (M.R.W.), the Whitehall Foundation (M.R.W.), and Cornell University.

## AUTHOR CONTRIBUTIONS

Conceptualization, B.J.S., R.J.P., D.A.B., and M.R.W.; Methodology, B.J.S., R.J.P., D.A.B., R.B.E., and M.R.W.; Software, B.J.S., D.A.B., and R.B.E.; Investigation, B.J.S., R.J.P., D.A.B., V.L., K.H. and M.R.W.; Analysis, B.J.S., Writing – Original Draft, B.J.S. and M.R.W.; Writing – Review & Editing, B.J.S., R.J.P., D.A.B., and M.R.W.; Supervision, M.R.W.

## DECLARATION OF INTERESTS

The authors declare no competing interests.

## STAR★Methods

### KEY RESOURCES TABLE

| REAGENT or RESOURCE | SOURCE | IDENTIFIER |
| --- | --- | --- |
| **Bacterial and Virus Strains** |  |  |
| AAV9-CAG-Flex-GCaMP6s | UPenn VectorCore | N/A |
| AAVDJ-hSyn-DIO-GCaMP6m | Stanford Gene Vector and Virus Core | GVVC-AAV-95 |
| AAV5-CAG-Flex-GCaMP6f | UPenn VectorCore | N/A |
| AAV9-CAG-Flex-GFP | UNC Vector Core | N/A |
| AAV5-CAG-Flex-GFP | UNC Vector Core | N/A |
| AAV9-EF1α-DIO-eNpHR3.0-eYFP | Vector Biolabs | N/A |
| **Chemicals** |  |  |
| DAPI | Sigma-Aldrich | D9542-5mg |
| DABCO | Sigma-Aldrich | 290734-100ML |
| **Experimental Models: Organisms/Strains** |  |  |
| Mouse: Tg(Klk8-cre)NP171Gsat/Mmucd | GENSAT | RRID: MMRRC_036080-UCD |
| Mouse: B6.SJL-Slc6a3 tm1.1(cre)Bkmn/J | The Jackson Laboratory | RRID: IMSR_JAX:006660 |
| Mouse: STOCK Slc17a6 tm2(cre)Lowl/J | The Jackson Laboratory | RRID: IMSR_JAX:016963 |
| Mouse: C57BL/6J | The Jackson Laboratory | RRID: IMSR_JAX:000664 |
| **Software and Algorithms** |  |  |
| MATLAB script for analysis | Brianna J. Sleezer | N/A |
| Med Associates script for operant conditioning | Dave A. Bulkin, Ryan J. Post | N/A |
| MATLAB toolbox for multi-unit electrophysiology processing | Daniel N. Hill, Samar B. Mehta, David Kleinfeld | N/A |

### RESOURCE AVAILABILITY

#### Lead Contact

Further information and requests for resources and reagents should be directed to the Lead Contact, Melissa R. Warden.

### Materials Availability

This study did not generate new reagents.

### Data and Code Availability

Data and source code supporting the current study will be made available upon request.

## EXPERIMENTAL MODEL AND SUBJECT DETAILS

### Animals

All experiments were carried out under protocols approved by Cornell University’s Institutional Animal Care and Use Committee and conformed to NIH guidelines. Both male and female mice (postnatal 3-6 months) were used in this study. Mice were group housed in a vivarium on a standard 12h light/dark cycle. All experiments were conducted during the dark portion of the cycle. Klk8-Cre (GENSAT, RRID: MMRRC_036080-UCD, New York, NY), DAT-Cre (The Jackson Laboratory, Bar Harbor, ME), and Vglut2-Cre (The Jackson Laboratory, Bar Harbor, ME) mice were used for photometry experiments. Klk8-Cre mice were used for optogenetic inhibition experiments. C57BL/6J mice were used for electrophysiology experiments. All Cre driver lines were fully backcrossed to C57BL/6J mice. Mice were provided with ad libitum access to food and water prior to training on the poke-reward task.

## METHOD DETAILS

### Viral Vectors

For photometry experiments, Klk8-Cre (NP171) and Vglut2-Cre mice were injected with AAV9-CAG-Flex-GCaMP6s (Penn Vector Core, Philadelphia, PA). DAT-Cre mice were injected with AAV5-Syn-Flex-GCaMP6f (Penn Vector Core, Philadelphia, PA) or AAV-DJ-EF1α-DIO-GCaMP6m (Stanford Vector Core, Stanford, CA). For optogenetic experiments, Klk8-Cre (NP171) mice were injected with AAV9-EF1α-DIO-eNpHR3.0-eYFP (Vector Biolabs, Malvern, PA). For all experiments, control animals were injected with AAV9-CAG-Flex-GFP (UNC Vector Core, Chapel Hill, NC).

### Surgical Procedures

Mice were anesthetized with isoflurane (5%). Fur was trimmed, and mice were placed in a stereotaxic frame (Kopf Instruments, Tujunga, CA) on a heating pad to prevent hypothermia. Isoflurane was delivered at 1-3% throughout surgery; this level was adjusted to maintain a constant surgical plane. Ophthalmic ointment was used to protect the eyes. Lactated ringers (500 ml, subcutaneous) was administered before the start of surgery. A mixture of 0.5% lidocaine and 0.25% bupivicaine (100 ml) was injected subdermally along the incision line. The scalp was disinfected with betadine and alcohol. A midline incision exposed the skull, which was thoroughly cleaned, and a craniotomy was made above the LHb or VTA. Virus was targeted to the LHb (−1.80 AP, 0.40 ML, −2.80 and −2.60 DV) or VTA (−3.10 AP, 0.35 ML, −4.60 and −4.30 DV), and slowly pressure injected (100 nl min^-1^) using a 10 ml Hamilton syringe (nanofil; WPI, Sarasota, FL), a 33 gauge beveled needle, and a micro-syringe pump controller (Micro 4; WPI, Sarasota, FL). After each injection, the needle was left in place for 10 minutes and then slowly withdrawn. For photometry experiments, a total of 600 nl (300 nl at each DV site) of vector was injected and an optical fiber (400 µm diameter, 0.48 NA, Doric Lenses, Québec, Canada) was implanted in the right hemisphere. Implants were targeted to LHb (−1.70 AP, 0.40 ML, −2.40 DV) or VTA (−3.10 AP, 0.35 ML, −4.45 DV). For LHb inhibition experiments, a total of 1200 nl (300 nl at each DV site in each hemisphere) of vector was injected and an optical fiber (200 µm diameter, 0.22 NA, Thorlabs Inc., Newton, NJ) was implanted at a 20 degree angle in each hemisphere (−1.80 AP, +-1.23 ML, −2.21 DV). A layer of metabond (Parkell, Inc., Edgewood, NY) and dental acrylic (Lang Dental Manufacturing, Wheeling, IL) was applied to firmly hold the fiber in place, and the surrounding skin was sutured closed. Post-operative buprenorphine (0.05 mg/kg) and carprofen (5 mg/kg) were administered subcutaneously. Virus was allowed to express for a minimum of 6 weeks before behavioral testing.

### Immunohistochemistry

To determine the specificity of the Klk8-Cre (NP171) to the LHb, male Klk8-Cre (NP171) mice were injected with AAV9-CAG-Flex-GFP in the LHb, as described above. After 4 weeks of expression, mice were perfused with PBS and 4% paraformaldehyde. Brains were extracted, post-fixed in 4% PFA for 24 h, and stored in 30% sucrose in PBS until sliced into 50 µm coronal sections. Sections containing the LHb were blocked (10% normal goat serum; ThermoFisher Scientific, Waltham, MA) and incubated with 1:400 anti-NeuN primary antibody (ABN78, EMD Millipore, Darmstadt, DE) and 1:500 Cy3-expressing secondary antibody (AB_2307443, Jackson Immunoresearch, West Grove, PA) (Zhong et al., 2017). Sections were imaged under a confocal microscope (LSM800, Zeiss, White Plains, NY) to determine the specificity of the NP171 marker for neurons and the penetrance of Cre-dependent viral vectors for neurons in the injection area.

### Fiber Photometry

Fiber photometry was performed using a Doric photometry system. A 490 nm LED was sinusoidally modulated at 211 Hz and passed through a GFP excitation filter. A 405 nm LED was modulated at 531 Hz and passed through a 405 nm bandpass filter. Both light streams were coupled to an optical fiber patch cord (0.48 NA; 400 μm core), which was connected to an optical fiber brain implant in each mouse. GCaMP6 fluorescence was collected by the same fiber, passed through a GFP emission filter, and focused onto a photoreceiver. To calculate ΔF/F, a least-squares linear fit was applied to the 405 nm signal to align it to the 490 nm signal, producing a fitted 405 nm signal that was used to normalize the 490 nm according to the following: ΔF/F = (490 nm signal − fitted 405 nm signal)/fitted 405 nm signal.

### Multi-Unit Electrophysiology

To record multi-unit activity (MUA), we implanted male C57BL/6 mice with 16-channel electrode arrays (35 µm tungsten electrodes with 200 µm electrode spacing, 200 µm row spacing, 6 mm in length; Innovative Neurophysiology, Inc., Durham, NC) centered over the right LHb. Target coordinates, relative to bregma, were: AP: −1.20 mm to −2.80 mm (anterior and posterior edges of the array); ML: 0.30 mm and 0.50 mm (medial and lateral electrode rows, respectively); DV: − 2.55 mm. A ground wire, affixed to the array, was attached to two stainless steel screws placed in the cerebellum. We used a Tucker-Davis Technologies acquisition system and Synapse software to record spike data (Tucker-Davis Technologies, Alachua, FL). Voltage measurements were collected and saved at 24.414 kHz. MUA was extracted from the raw voltage trace through a series of offline processing steps. First, signals were filtered and large artifactual voltage fluctuations were removed from each channel using stationary wavelet decomposition/transform. Next, a common average reference for each array was calculated by taking the sample by sample average of all channels; this global average was then subtracted from the signal on each channel. This method of referencing has been found to outperform alternative referencing methods, such as single best electrode referencing (Ludwig et al., 2009). MUA spiking activity was then extracted using the toolbox UltraMegaSort2000 and a voltage threshold of 2.5 standard deviations above the mean of the voltage trace. To determine the location of electrode tips, we used an optical clearing technique that allowed us to visualize the entire electrode tract (including electrode tips) for all 16 electrodes in each array. Following collection of MUA data, mice were perfused using saline and 4% paraformaldehyde and decapitated. Skin and other tissues were removed from the head (keeping electrode arrays intact) and skulls (with intact arrays) were drop fixed in 4% paraformaldehyde for 48 to 72 hours at 4°C. Brains were then carefully dissected from the skull and arrays were gently removed. A 1.5 mm thick sagittal slice of brain tissue centered around the electrode array was taken from each brain. Slices were then drop fixed in glutaraldehyde for 24 hours at 4°C, washed in PBS-T for 24 hours, and then placed in 6% SDS at 37°C and checked daily to monitor clearing progress. Slices were typically sufficiently cleared in 6 to 10 days. Slices were then washed in PBS-T at 37°C for 48 hours and subsequently placed in an iodixanol solution composed of 50 g diatrizoic acid, 40 g N-methyl-d-glucamine, 55 g iodixanol, and 0.02% sodium azide per 100 ml water (Murray et al., 2015). Slices were gently swirled daily and monitored for transparency and refraction changes over the course of 2 to 5 days. Samples were then transferred to a fresh iodixanol solution for 24 hours and, finally, mounted in the iodixanol solution between two cover glasses which were separated by 1.5 mm rubber gaskets. Slices were then imaged using a confocal microscope (LSM800, Zeiss, White Plains, NY). A 647 nm wavelength excitation light was used for imaging. To determine the location of the medial and lateral electrode rows, z-stacks were constructed from optical sections taken in 50 µm increments. Given that electrode arrays extended posterior to LHb, the posterior most 3 to 4 electrodes in both the medial and lateral rows were typically excluded from further analysis based on inspection of the electrode tract locations.

### Optogenetic Inhibition

During behavioral testing, external patch cords (200 µm diameter, 0.22 NA, Doric Lenses, Québec, Canada) were coupled to implanted fiber optic cannulae (CFM22U-20, Thor Labs, NJ, US) with zirconia sleeves. Cannulae were placed above LHb bilaterally as described above. An optical commutator allowed for unrestricted rotation (Doric Lenses, Québec, Canada). Optical inhibition was provided with 594 nm laser light diode pumped solid state laser (Mambo 100, Colbalt, Solna, SE). Inhibition experiments used 4 mW light (127 mW/mm^2^ at the fiber tip).

### Behavioral Testing

In all versions of the poke-reward task, mice were first water restricted over the course of three to five days until they reached 80% of their pre-restriction body weight. After restriction, mice were trained to poke their nose into a hole (ENV-313W; Med Associates, VT, US) on one wall of a 20 x 22 cm operant chamber (ENV-307W-CT) housed in a sound-attenuating box (ENV-022MD) and containing a lickometer (ENV-250B) on the end of the chamber opposite the nose-poke. At the start of each session, a light in the nose-poke hole was illuminated. Upon completion of a successful nose-poke, the light in the nose-poke hole was turned off, a soft white noise was turned on to indicate the availability of reward, and a water reward was delivered via syringe pump (PHM100) to a fluid port on the opposite wall of the chamber. A nose-poke entry followed by a lick was considered a single trial. Mice were free to run back and forth between the nose-poke hole and the reward spout and complete trials at their own pace. Training continued daily until mice were able to perform 90 or more trials within a 30 minute session across two consecutive daily sessions. Following training, mice were placed on other versions of the task. Mice were run on one 30-minute session each day, 7 days a week.

#### Probabilistic task

In the probabilistic version of the poke-reward task, each successful nose-poke had a 20% chance of yielding a large reward (20 µl of water), a 60% chance of yielding a medium reward (10 µl of water) and a 20% chance of yielding no reward (0 µl of water). No cues were provided to indicate the reward size or probabilities.

#### Blocked task

In the blocked version of the poke-reward task, no reward trials (0 µl of water) and reward trials (10 µl of water) were grouped into 5-minute long blocks of trials that alternated between reward available and no reward available. Because we wanted to be sure that mice experienced each block, block reward contingencies were only changed once animals completed a successful nose-poke following 5 minutes within a block − this design minimized instances in which mice may not complete any trials during a 5-minute long period and subsequently would not be aware of the block transition. As such, the absolute duration of each block was a minimum of 5 minutes, but varied depending on mouse behavior. All photometry experiments for the poke-reward task were run using the same mice and occurred in the following order: training (5-10 days), probabilistic task (5 days), blocked task (5 days).

#### Optogenetic inhibition: Re-engagement task

For optogenetic experiments only, mice performed a task designed to probe the capacity for LHb inhibition to drive task re-engagement after entering a disengaged state (**Figure 6C**). In this task, we provided mice with a medium reward (10 µl water) following each successful nose poke, consistent with the training paradigm above. We then waited for mice to disengage from the task (we defined disengagement as the point at which mice had not completed a trial - i.e., a successful nose poke - over the course of a 2-minute period). We then turned the laser on (or began a sham laser off period), and measured the latency for mice to re-engage (i.e. perform a successful nose poke) in the task. We collected behavioral data over the course of four consecutive daily sessions, alternating between sessions in which the laser was turned on and sessions in which the laser was off. The order of laser on and laser off sessions was counterbalanced across mice. Following the completion of these four sessions, mice performed an additional four sessions on a similar version of the re-engagement task. In these sessions, all task and laser conditions were the same, with the exception that all trials yielded no reward (0 µl).

#### Optogenetic inhibition

*Persistence task*. For optogenetic experiments only, mice were run on a task that was designed to probe the capacity for LHb inhibition to drive task persistence (**Figure 6E**). In this task, we again provided mice with a medium reward (10 µl water) following each successful nose poke, consistent with the training paradigm above and waited for mice to disengage from the task (once again, we defined disengagement as the point at which mice had not completed a successful nose poke over the course of a 2-minute period). In this version of the task, we waited for mice to re-engage in the task (i.e. complete a successful nose poke) and then turned the laser on (or began a sham laser off period). We terminated the session once mice disengaged from the task once again. We measured persistence as the number of trials (successful nose pokes) mice performed during the laser on (or sham laser) period. We again collected behavioral data over the course of four consecutive daily sessions, alternating between sessions in which the laser was turned on and sessions in which the laser was off. The order of laser on and laser off sessions was counterbalanced across mice. Following the completion of these four sessions, mice performed an additional four sessions on a similar version of the persistence task. In these sessions, all task and laser conditions were the same, with the exception that all trials yielded no reward (0 µl).

### QUANTIFICATION AND STATISTICAL ANALYSIS

#### Identification of State Transitions (Hidden Markov Model)

To identify periods of high and low engagement, we modeled high and low engagement as two latent states that could underlie behavior. Because the level of engagement could be measured in multiple ways (the frequency of licks and/or the frequency of nose pokes), we used a multivariate HMM to simultaneously account for both types of observations. Behavior was coded as binary vectors indicating the presence or absence of each behavior during each second of the task. These emissions were modelled as Poisson random variables whose probability of occurring was free to differ across latent states. The model was fit via expectation-maximization using the Baum Welch algorithm (Bilmes, 1998; Murphy, 2012), which finds a (possibly local) maxima of the complete-data likelihood. The behavioral data was oversampled relative to the slow changes in engagement of interest here, so we used two methods to highlight the slow changes in engagement states. First, observations were smoothed across neighboring time bins (10 bins) to disrupt the irrelevant local structure that occurred because licks and nose pokes happened at opposite ends of the chamber. Second, we added a regularization term that penalized frequent transitions between states (Montanez et al., 2015). The algorithm was initialized with a random seed once, and the model that maximized the observed (incomplete) data log likelihood was ultimately taken as the best for each session. Finally, we used the Viterbi algorithm to discover the most probable a posteriori sequence of latent states, given the model and behavioral observations (Murphy, 2012).

### Statistics

All statistical analyses were performed using MATLAB (MathWorks, Natick, MA). Effects with a P value less than 0.05 were considered significant. All variance estimates and error bars represent standard error of the mean (SEM).

## Supplemental Information

**Figure S1 (Related to Figure 1).**
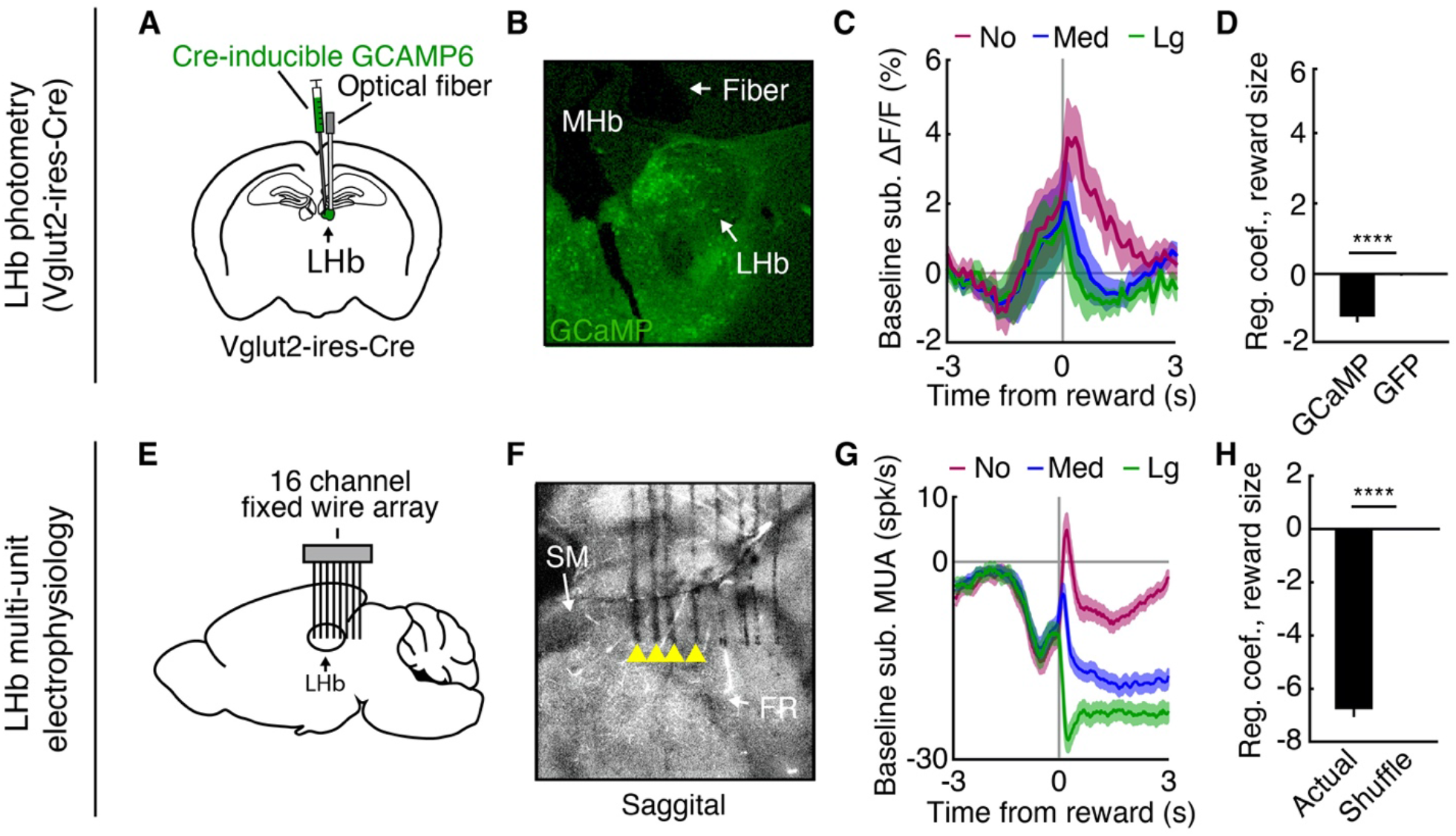
Phasic LHb Activity Upon Reward Omission, Vglut2-Cre Photometry and Multi-unit Electrophysiology. (A) Schematic of LHb viral vector injection and optical fiber placement, Vglut2-Cre mice. (B) LHb GCaMP6 expression, Vglut2-Cre mouse. (C) Average baseline-subtracted ΔF/F in Vglut2-Cre mice (n = 3) during probabilistic task sessions, aligned to reward receipt and separated by reward size (no reward, red; medium reward, blue; large reward, green). (D) Average regression coefficients obtained by regressing ΔF/F 0-1 s following the start of reward consumption onto reward size in GCaMP (n = 3) and GFP (n = 3) Vglut2-Cre mice. ****p < 0.0001, two-sample t-test. (E-H) Same as in A-D, but for LHb multi-unit electrophysiology (n = 40 electrodes in 6 mice). ****p < 0.0001, t-test compared to shuffled sample. Shaded regions and error bars indicate SEM.

**Figure S2 (Related to Figure 2).**
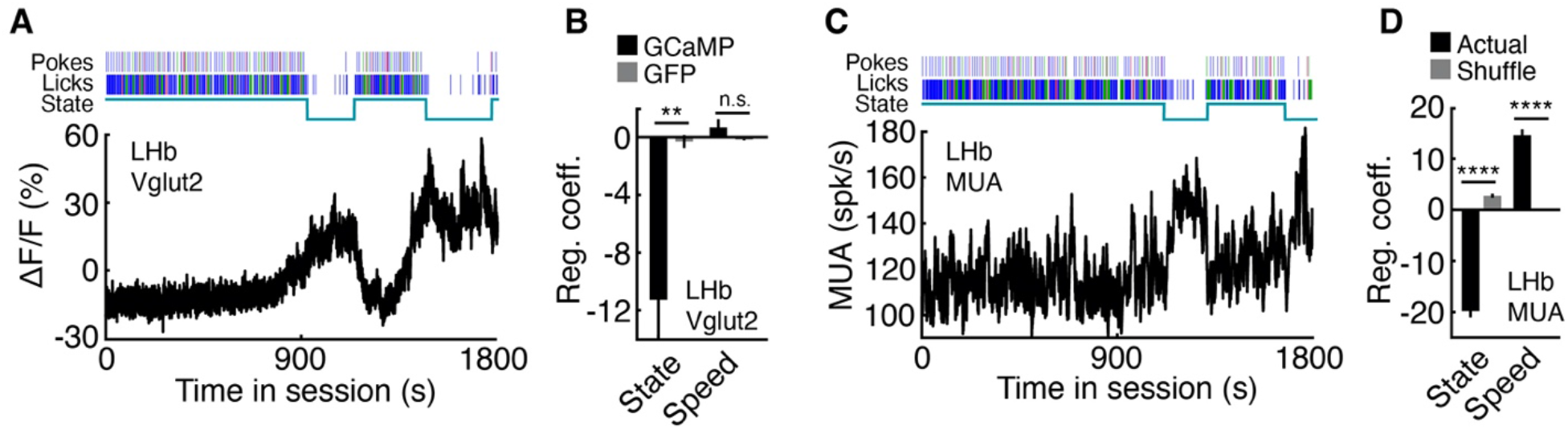
Tonically Elevated LHb Activity During Spontaneous End-of-Session Task Disengagement, Vglut2-Cre Photometry and Multi-unit Electrophysiology. (A) Example LHb Vglut2-Cre photometry during a complete probabilistic task session. Lines above the plot indicate all poke and lick times (no reward, red; medium reward, blue). The line under the poke and lick timestamps indicates whether mice were in a high or low task engagement state. (B) Average regression coefficients obtained by regressing ΔF/F onto task engagement state and running speed in GCaMP (n = 3) and GFP (n = 3) Vglut2-Cre mice. **p < 0.01, two-sample t-test. (C) Example LHb multi-unit activity during a complete probabilistic task session. Same conventions as A. (D) Average regression coefficients obtained by regressing multi-unit activity onto engagement state and running speed (n = 40 electrodes in 6 mice). ****p < 0.0001, t-test compared to shuffled sample. Error bars indicate SEM.

**Figure S3 (Related to Figure 3).**
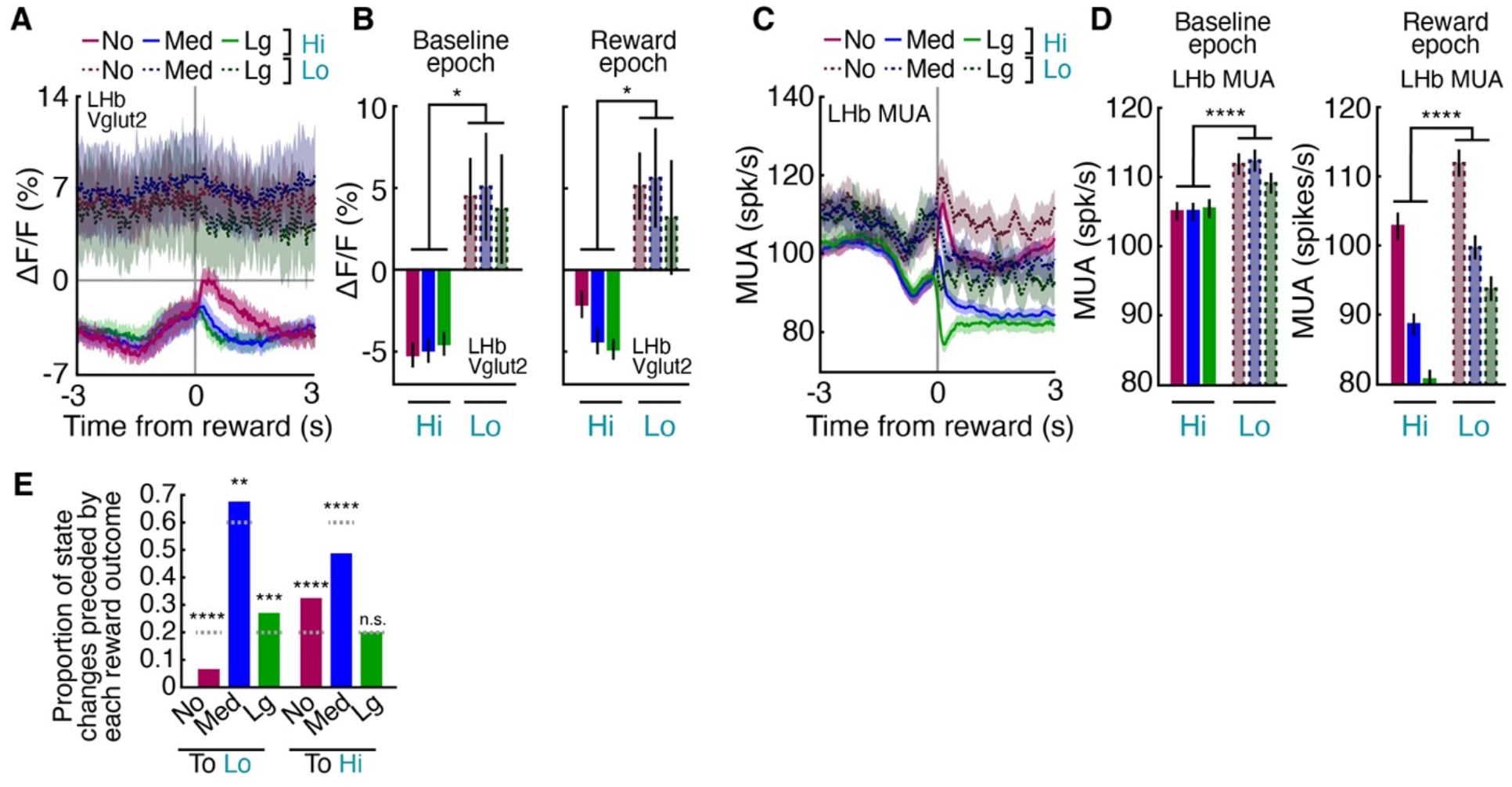
Baseline LHb Activity Reflects Task Engagement State, Vglut2-Cre Photometry and Multi-unit Electrophysiology. (A) Average ΔF/F in GCaMP Vglut2-Cre mice (n = 3) during probabilistic task sessions, aligned to reward receipt and separated by engagement state (solid versus dotted lines) and reward size (no reward, red line; medium reward, blue; large reward, green). (B) Average ΔF/F during baseline and reward epochs in high and low engagement states. *p < 0.05, paired samples t-test. (C) Average multi-unit activity (n = 40 electrodes in 6 mice) during probabilistic task sessions, aligned to reward receipt and separated by reward size. Same conventions as A. (D) Average multi-unit activity during baseline and reward epochs in high and low engagement states. ****p < 0.0001, paired samples t-test. (E) Proportion of transitions to low and high engagement states preceded by trials that yielded no reward, medium reward, or large reward. Chance proportions indicated by dashed gray lines. **p < 0.01, ***p < 0.001, ****p < 0.0001, binomial test. Shaded regions and error bars indicate SEM.

**Figure S4 (Related to Figure 4).**
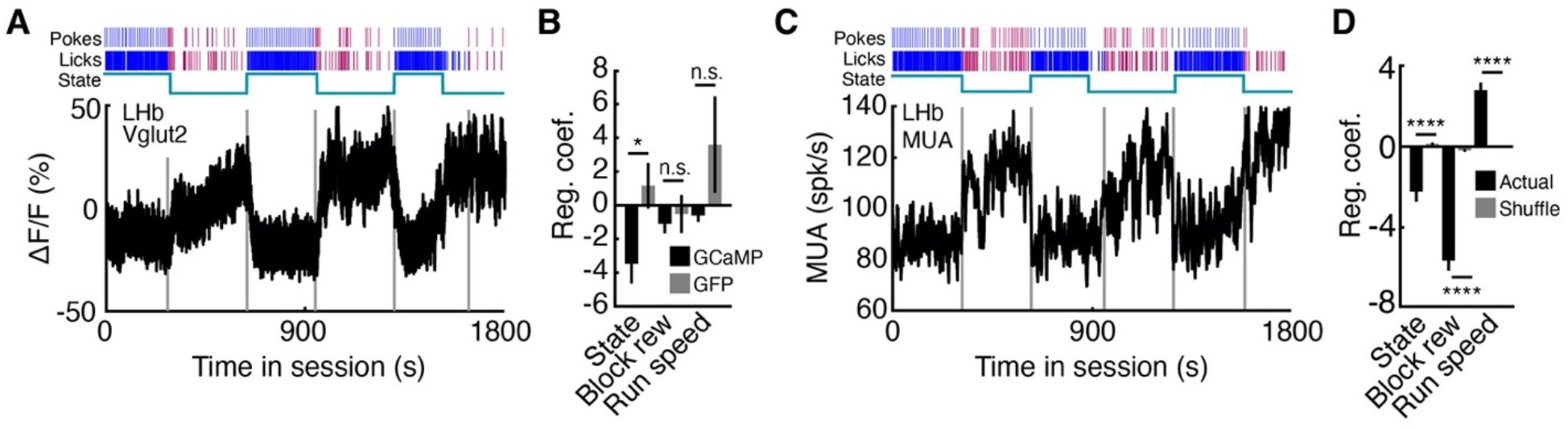
LHb Activity is Tonically Elevated During Task Disengagement Due to Repeated Reward Omission, Vglut2-Cre Photometry and Multi-unit Electrophysiology. (A) Example LHb Vglut2-Cre photometry during a blocked task session. Lines above the plot indicate all poke and lick times (no reward, red; medium reward, blue). The line under the poke and lick timestamps indicates whether mice were in a high or low task engagement state. (B) Average regression coefficients obtained by regressing ΔF/F onto engagement state, block reward, and running speed in GCaMP (n = 3) and GFP (n = 3) Vglut2-Cre mice. *p < 0.05, two-sample t-test. (C) Example LHb multi-unit spiking activity during a blocked task session. Same conventions as A. (D) Average regression coefficients obtained by regressing multi-unit activity onto engagement state, block reward, and running speed (n = 40 electrodes in 6 mice). ****p < 0.0001, t-test compared to shuffled sample. Error bars indicate SEM.

**Figure S5 (Related to Figure 6).**
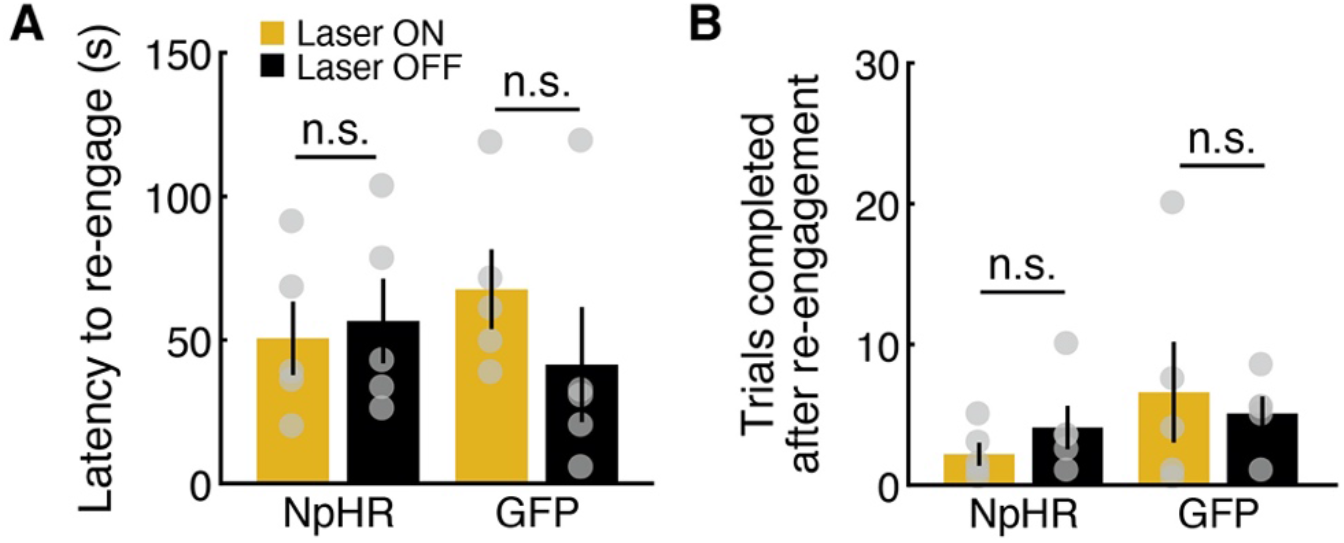
LHb Inhibition Does Not Extend Reward-Seeking Behavioral States When Rewards Are Not Available. (A) Average latency to re-engage in task performance without rewards. LHb inhibition initiated during disengagement. NpHR (n = 5) and GFP (n = 5) Klk8-Cre (NP171) mice. (B) Average number of trials completed after task re-engagement without rewards. LHb inhibition initiated upon re-engagement. NpHR (n = 5) and GFP (n = 5) Klk8-Cre (NP171) mice. Paired samples t-test. Error bars indicate SEM.

## Notes

### Competing Interest Statement

The authors have declared no competing interest.

